# SimpactCyan 1.0: An Open-source Simulator for Individual-Based Models in HIV Epidemiology with R and Python Interfaces

**DOI:** 10.1101/440834

**Authors:** Jori Liesenborgs, Diana M Hendrickx, Elise Kuylen, David Niyukuri, Niel Hens, Wim Delva

## Abstract

SimpactCyan is an open-source simulator for individual-based models in HIV epidemiology. Its core algorithm is written in C++ for computational efficiency, while the R and Python interfaces aim to make the tool accessible to the fast-growing community of R and Python users. Transmission, treatment and prevention of HIV infections in dynamic sexual networks are simulated by discrete events. A generic “intervention” event allows model parameters to be changed over time, and can be used to model medical and behavioural HIV prevention programmes. First, we describe a more efficient variant of the modified Next Reaction Method that drives our continuous-time simulator. Next, we outline key built-in features and assumptions of individual-based models formulated in SimpactCyan, and provide code snippets for how to formulate, execute and analyse models in SimpactCyan through its R and Python interfaces. Lastly, we give two examples of applications in HIV epidemiology: the first demonstrates how the software can be used to estimate the impact of progressive changes to the eligibility criteria for HIV treatment on HIV incidence. The second example illustrates the use of SimpactCyan as a data-generating tool for assessing the performance of a phylodynamic inference framework.

## Introduction

Mathematical models are a commonly used to study key features of infectious disease epidemics. In this way, past or future dynamics of disease progression, transmission, prevention and treatment can be simulated. Deterministic compartmental models are the most popular. Using systems of difference or differential equations, these models represent how the size of population subgroups (e.g. stratified by infection status and disease stages) change over time. However, the implicit assumption of homogeneity of individuals within the same compartment may be particularly inadequate for sexually transmitted diseases, as the behavioural and biological drivers of these diseases are subject to high individual heterogeneity. Rather than simulating population averages, individual-based models (IBMs) simulate events that happen to specific individuals. Because of this, they are able to take various sources of individual heterogeneity into account^1^.

IBMs allow for a bottom-up approach to modelling complex systems: population-level features are not modelled directly, but emerge from processes and events that are modelled at the level of the individual and its immediate environments. Thanks to multi-core processors and increased access to high-performance computing infrastructure, the computational cost of IBMs has become less prohibitive. As a result, a growing use of IBMs in infectious disease epidemiology has been observed^2^.

A large amount of general frameworks for individual-based simulations have been developed in the last decades. These platforms vary widely in terms of platform properties, usability, operating ability, pragmatics and security management, making it difficult to choose the most suitable framework for simulation in the context of a particular research question^3^.

Table 1 summarises functional and structural differences between SimpactCyan and ten other tools for constructing individual-based models of HIV transmission between individuals connected via sexual relationships. These tools were identified by an ongoing systematic review of simulation-based methods for the calibration of individual-based models to summary data in epidemiology^4^.

**Table 1.** Functional and structural differences between SimpactCyan and existing tools for IBM studies in HIV epidemiology.

| Tool | Time imple-<br>mentation | Sexual net-<br>work | R interface | Python inter-<br>face | Source code<br>available<br>online | Reference |
| --- | --- | --- | --- | --- | --- | --- |
| SimpactCyan | continuous,<br>mNRM | dynamic | ✓ | ✓ | ✓ | 5–7 |
| GEMFsim | continuous,<br>Gillespie<br>algorithm | static | ✓ | ✓ | ✓ | 8 |
| FAVITES | continuous,<br>Gillespie<br>algorithm | static | ✓ | ✓ | ✓ | 9 |
| EpiModel | discrete | dynamic | ✓ | ✗ | ✓ | 10 |
| STDsim | discrete | dynamic | ✗ | ✗ | ✗ | 11 |
| NetLogo | discrete | dynamic | ✓ | ✗ | ✓ | 12, 13 |
| EMOD | discrete | dynamic | ✗ | ✓ | ✓ | 14 |
| HIV-CDM | discrete | dynamic | ✗ | ✗ | ✗ | 15 |
| MicroCOSM | discrete | dynamic | ✗ | ✗ | ✗ | 16 |
| Path 2.0 | discrete | dynamic | ✓ (through<br>RNetLogo) | ✗ | ✗ | 17 |
| TITAN | discrete | dynamic | ✓ | ✓ | ✓ | 18 |

As evident from Table 1, available tools for IBM studies in HIV epidemiology vary in their features and limitations. Some tools simulate static instead of dynamic sexual networks. For others the source code is not freely available, or interfaces for Python or R (the most popular languages among infectious disease modellers) do not exist. Run time can be a limitation as well. NetLogo, for example, is a wonderful tool for building simple, didactic models^3^. The problem is that they become painstakingly slow when HIV epidemics are simulated for more than a few hundred time steps and in populations larger than a few hundred individuals.

With a few exceptions (the GEMFsim^8^ and FAVITES^9^ simulators), existing simulators that can be readily used in HIV epidemiology implement IBMs in discrete time rather than continuous time. By simulation in continuous time we mean that events can take place and subsequently the state of the system can be updated at any point in time. It also means that the time interval between the execution of two consecutive events is only limited by the numeric precision of the implementation of the method used to sample event times. The advantage of a continuous time implementation of IBMs is the elegance with which it can handle competing risks to multiple events. For instance, an individual may be at risk of infecting his partner with HIV while also being at risk of dying because of AIDS. Fixed time steps can result in a situation where both these events are scheduled to take place between now and the next time step. However, this can only happen if the transmission event happens first. In contrast, evaluating the model in continuous time means the method is explicit about the order in which the two events are scheduled. Whichever event that is, the execution of that event includes processing the logical consequences for the likelihood of other, subsequent events. Furthermore, evaluation in continuous time means that the event scheduler “jumps” from one event to the next. In contrast, when the model is evaluated in fixed time steps, simulations that involve both common and rare events can be computationally inefficient. Commonly occurring events will require a small time step, which may lead to the occurrence of rare events being evaluated at a higher frequency than necessary.

To our knowledge, all existing implementations of IBMs for dynamic sexual networks consider the event of relationship formation in a sequential manner. As a consequence they require ad-hoc assumptions and decisions about in what order people “go out” to find partners and can “be found”. For example, in EMOD, males and females are placed in a separate queue, where they stay for a predefined period, after which they form a relationship^14^. In STDsim, the order of going out to search for a partner and for being available for a relationship are both determined by stochastic processes^11^.

SimpactCyan was conceived as a simulator for building IBMs to address research questions in HIV epidemiology. As we show in this paper, IBMs built with SimpactCyan can be used to interrogate hypotheses in the spheres of network and social epidemiology, computational biology, public health policy. The simulator is evaluated in continuous time. Each time an event happens, the state of the system is updated. Furthermore, the time-evolution of the sexual network is modelled by considering all possible relationships simultaneously instead of sequentially.

Simpact (*SimpactWhite*) was first developed in Matlab^19–21^. Later, variants were developed as a MASON Multi-agent Simulation Toolkit in Java (*SimpactBlue*), and in Python (*SimpactyPurple*)^22^. To improve both speed and user-friendliness of the tool, we embarked on a major overhaul in 2013, leading to the current version (*SimpactCyan*) that combines a computationally efficient simulation engine written in C++ with R and Python interfaces. An exhaustive, deep comparison of SimpactCyan with all prior Simpact programs is beyond the scope of this paper, and arguably not crucially important, for the simple reason that there is no ongoing development of nor support for any of the legacy versions (SimpactWhite, SimpactBlue and SimpactPurple) and these versions are no longer in use. However, it may be useful to give some perspective of the relative improvement. In the early stages of SimpactCyan development, we conducted a comparison study that indicated runtimes for SimpactCyan were up to 280 times shorter, compared to SimpactWhite^23^.

In this paper, we describe a more efficient variant of the modified Next Reaction Method (mNRM) that drives the simulator, we outline key built-in features and assumptions of individual-based models formulated in SimpactCyan, and provide code snippets for how to formulate, execute and analyse models in SimpactCyan through its R and Python interfaces. The complete documentation for SimpactCyan is availabe at https://simpactcyan.readthedocs.io/en/latest/. As runtimes for SimpactCyan strongly depend on population size and the intensity with which relationships are formed and dissolved in the population, we present the results of a concise simulation study to provide additional insights and visual representation of these associations. We end by giving two examples of applications in HIV epidemiology: the first demonstrates how the software can be used to estimate the impact that changes to the eligibility criteria for antiretroviral therapy (ART) had on HIV incidence in a hyperendemic setting. The second example illustrates how SimpactCyan can be used to generate data for evaluating the performance of other frameworks for modelling HIV transmission dynamics.

## Discrete events simulation algorithm

### The modified Next Reaction Method (mNRM)

Event times, i.e. time points in the simulation at which events are scheduled to take place, are determined using the *modified Next Reaction Method* (mNRM)^24^, a more efficient variant of the Gillespie algorithm^25–27^ and the Next Reaction Method^28^. The mNRM was originally designed for simulating chemical systems with time-dependent propensities and delays, but in SimpactCyan we use it to simulate how individuals are at risk of events according to time-dependent hazard functions^29^. In the mNRM algorithm, there is a core distinction between *internal event times* and (simulated) *real-world event times*. The internal event times determine when an event will be triggered according to the event’s *internal clock*. Internal clock time advances faster as the hazard for the event increases. By real-world time we mean the calendar time in the simulated population.

Calling the *internal* time interval until a specific event fires Δ*T*, such internal time intervals are randomly sampled from an exponential distribution: Δ*T* ∼ Exp(1).

The event’s *hazard function h*(•), referred to as the *propensity function* in^24^, maps the internal time interval Δ*T* until the event fires onto Δ*t*, a real-world time interval,

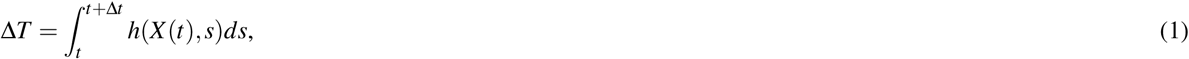

in which *t* is the previous time an event was triggered. (This corresponds to equation 13 in the original article^24^, where we have omitted the event-specific index for concision.) It is this hazard *h* that can depend on the state *X* (*t*) of the simulation, and possibly also explicitly on time *t*. In SimpactCyan, the state of the simulation is made up of all the individuals in the population and their respective properties, such as their age, gender, HIV infection status, ART status, and whom they are in relationships with. This state *X* (*t*) does *not* depend on time in a *continuous* manner, it only changes when an event is fired, i.e. when its internal time interval expires. Note that the formula above is for a single event, and while Δ*T* itself is not affected by other events, the mapping onto Δ*t* certainly can be: other events can change the simulation state, and the hazard of the event depends on this state.

The main idea is illustrated in Figure 1: internal time intervals are chosen from an exponential distribution, and are mapped onto real-world time intervals through hazard functions. Because hazards can depend on the simulation state and can have an explicit time dependency, this mapping can be rather complex.

**Figure 1.**
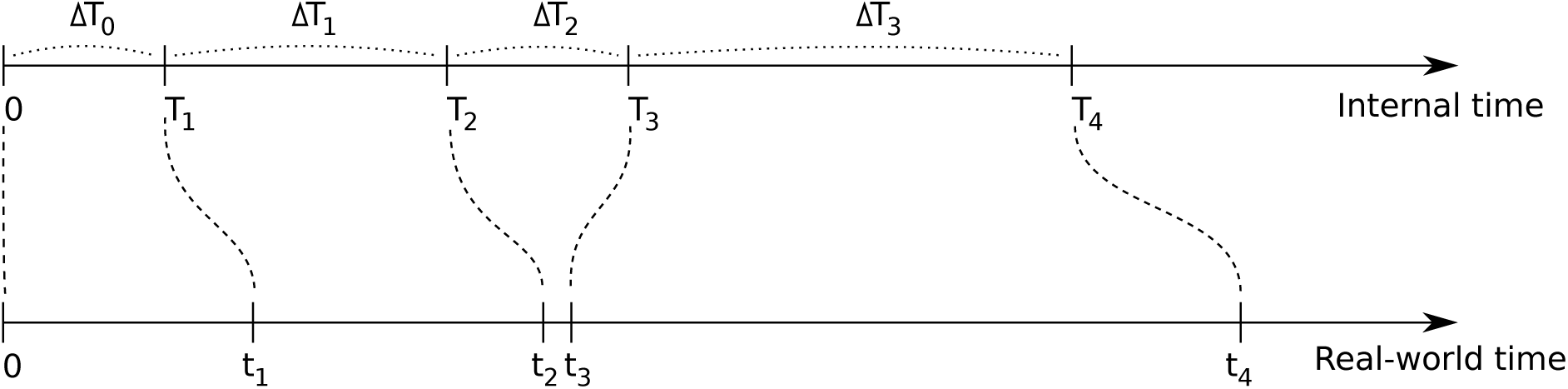
In the modified Next Reaction Method, intervals Δ*T*_*i*_ (in general for different events) are generated independently from other events in a straightforward manner, using an exponential probability distribution (Δ*T*_*i*_ ∼ Exp(1)), and are used to advance an *internal* clock *T*. Using the notion of a hazard function (1), these internal time intervals are mapped onto intervals Δ*t*_*i*_, which advance a (simulated) *real-world* time *t*, and need not have a straightforward relation to the internal times: a small internal time difference can lead to a large real-world time difference and vice versa. It is through this hazard function that interdependencies between events can be introduced.

While the hazard *can* cause complex behaviour, this is of course not necessarily the case. If one uses a constant hazard, this merely causes a linear scaling between internal time Δ*T* and real-world time Δ*t*:

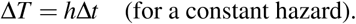

This also illustrates that the larger the hazard, the earlier the event will fire, i.e. the real-world time interval will be smaller.

As an example, let’s consider the event of forming a heterosexual relationship. At a certain time in the simulation, many formation events will be scheduled, one event for each man-woman pair that can possibly form a relationship. The internal time interval for each of these events will be drawn from the simple exponential distribution. The mapping to a real-world time at which the event will fire, is done using the hazard-based method, and the event that will take place next is the one that will have the smallest of these real-world times (cfr. the time ordering in step 6 of algorithm 3 in^24^). This hazard depends on aspects of the simulation state as defined by the hazard function for relationship formation: how many relationships the man and woman of the candidate couple are already engaged in, what the preferred age differences with their respective partners are, etc. One can also imagine an explicit time dependency in the hazard: e.g. the hazard of forming a relationship increases as the time period since the relationship became possible gets longer.

While most of the events in SimpactCyan are scheduled using the exponential distribution to generate values for internal Δ*T*, some events are scheduled directly in real-world time. An example of this is the scheduling of the HIV ‘seeding’ event, i.e. the timing of introducing HIV into the population. This alternative method could still be thought of as a special case of internal and real-world time mapping. This is because if Δ*T* is set to the actual real-world time interval until the event fires, and the hazard is set to *h* = 1, internal and real-world time intervals match.

### More efficient mNRM algorithm

Each time an event is triggered, the state of the simulation changes. Because the hazard of any event can depend on this state, in the most general version of the mNRM algorithm, one would recalculate the real-world event times of all remaining events each time an event gets triggered: this ensures that the possibly changed state is taken into account. Always recalculating all event fire times is computationally very inefficient, however. Although the state may have been changed somewhat, this change may not be relevant for many of the event hazards in use. As a result, most updated real-world event times would be the same as before.

To avoid unnecessary recalculations of event times, SimpactCyan employs a variant of the mNRM algorithm, in which each individual is linked to a list of events that involve him or her, and events that involve multiple people will appear on the lists of all of these individuals. For example, a mortality event would be present in the list of only one individual, while a relationship formation event concerns two people and would therefore appear on two such lists. Figure 2 illustrates this idea. The lists are meant to keep track of event times that may require recalculation as a result of another event firing. If the HIV-positive partner of Person X (not depicted in Figure 2) dies, this death will trigger several updates to the system, including that the deceased gets removed from the population, and dissolution events are triggered immediately (i.e. no recalculation needed) for all the relationships that the deceased was engaged in. Such dissolution events will subsequently lead to the removal of the HIV transmission event that was on Person X’s list (also without any recalculation required).

**Figure 2.**
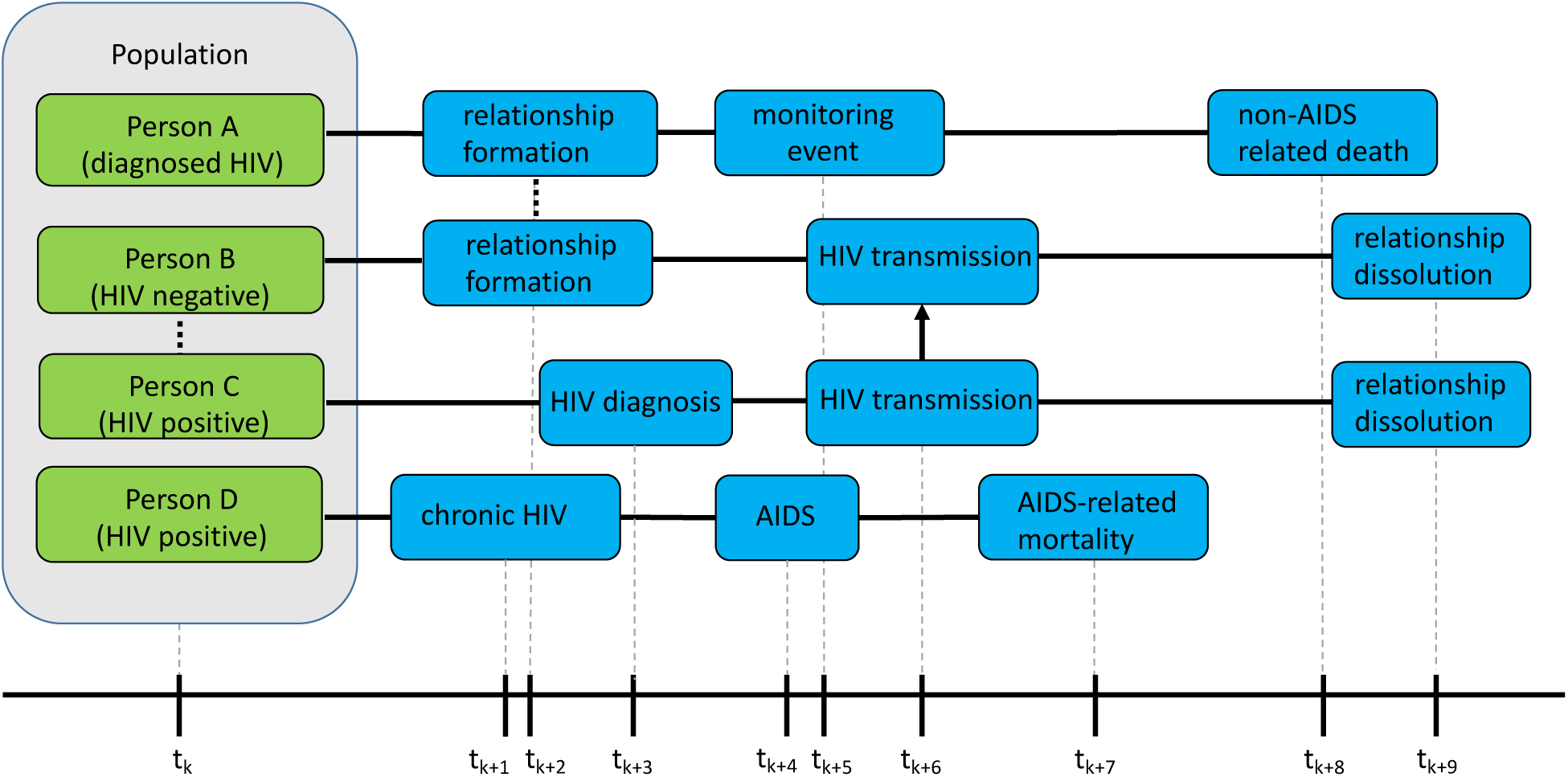
The Figure shows, for four different people in the population, what the next three scheduled events are after time point *t*_*k*_. Solid lines represent time intervals; the dotted line represents the formation of a relationship; the arrow represents HIV transmission. We can see that Person A and Person B will form a relationship. Concurrent with their relationship to Person A, Person B is also already in a relationship with Person C. We can see that an HIV transmission from Person C to Person B is scheduled, after Person C is diagnosed. After HIV transmission, Person B and Person C break up their relationship. After forming a relationship with Person B, Person A (who was already diagnosed with HIV) is monitored to follow up the progression of HIV. Person A dies a non-AIDS related death. Person D, who is HIV positive, will progress to the chronic stage of HIV infection, after which he or she will develop AIDS, and die of AIDS-related complications.

When an event fires, only the properties of a very limited set of people are changed, hence one only needs to recalculate the fire times of the events in those people’s lists. For example, when the event of Person A forming a relationship with Person B takes place, the real-world fire times for the events in the lists of Person A and Person B will be automatically recalculated. Apart from affecting the people in whose lists an event appears, some events can affect additional people. As an example, a birth event will only appear in the list of the pregnant woman and not in the event list of the father, because the scheduled birth should not be affected in the event of the death of the father. However, when triggered, the newborn will be listed as a child of the father. In general, the number of additionally affected people will be very small compared to the size of the population, causing only a fraction of the event fire times to be recalculated. This allows the modified algorithm to run much faster than the basic algorithm that always recalculates all event times. Furthermore, fire times of events that are present in the event lists of two individuals (e.g. relationship formation), are recalculated by only one of them.

Besides these types of events, there are also ‘global’ events. These events do not refer to a particular person and will modify the state in a very general way. In general, when such a global event is triggered, this causes *all* other event fire times to be recalculated. Introducing HIV into the population through an HIV seeding event is an example of a global event.

## Population and events in SimpactCyan

Model populations consist of men and/or women. They can be introduced into the simulation in two ways: (i) during the initialization of the simulation, in which case individuals with certain ages (drawn from a distribution) are added to the simulation, and (ii) through the birth of new individuals during the course of the simulation run.

Once born, an individual will become sexually active when a *debut event* is triggered. If the individual is introduced into the population at the start of the simulation, and the age exceeds the debut age, this event no longer needs to be scheduled. Every person always has a ‘normal’ *mortality event* scheduled, which corresponds to a cause of death other than AIDS.

To get an HIV epidemic started, an *HIV seeding event* must be scheduled. When this event is triggered, a number of people in the existing population will be marked as being HIV-infected. An infected individual will go through a number of infection stages, starting with acute HIV infection. After a default duration of 3 months^30^, a *chronic stage event* is triggered, moving the individual to the chronic infection stage. A fixed amount of time before dying of AIDS (15 months by default)^30^, an *AIDS stage event* is triggered, marking the transition of the chronic HIV stage to the AIDS stage. Six months before the expected AIDS-related death, a *final AIDS stage event* is triggered, after which the individual is in the ‘final AIDS stage’. It is assumed that one is too ill to be sexually active during this final stage^30^. When the *AIDS mortality event* is triggered, the individual dies of AIDS.

The `population.msm` parameter enables simulation of populations in which (some) men only form sexual relationships with other men, and/or can form relationships with with men and women. Under the default parameter setting (`population.msm = no`), every man-woman pair past the age of sexual debut can potentially form a relationship. For every such pair, a *formation event* is scheduled by sampling from the probability distribution that emerges as a result of the specified hazard function for relationship formation. An example for such a hazard function is the ‘agegap’ hazard, shown in the equation below. Only when an event of this type is triggered, an actual relationship is formed between the involved persons, which in turn can cause other events to get scheduled, e.g. a relationship dissolution event.

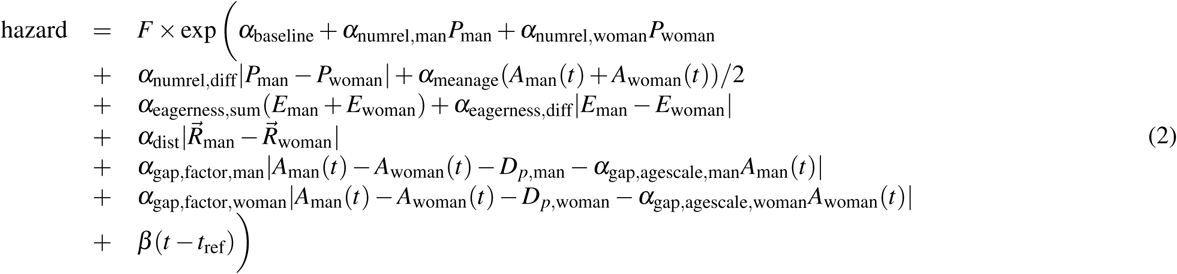

This complex looking hazard is actually of the form exp(*A* + *Bt*). The *α* parameters are weights that need to be set in the configuration of the simulation. They control the importance of various aspects of how individuals choose sexual partners. Via so-called intervention events, these weight parameters can be changed at arbitrary points in time during the simulation. In this way, temporal changes in sexual risk behaviours can be modelled, such as reductions in partner concurrency, age-disparate relationships or overall sexual activity levels. Variable *P* represents the number of relationships that an individual is currently engaged in, *A*(*t*) the age of the individual, and *E* represents a person-specific sex drive, of which the distribution is user-defined to allow control over the amount of heterogeneity in sexual activity within the population. The effect of distance 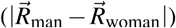 between the candidate partners can also be taken into account.

In the terms about the age gap, the age difference between the man and woman in the potential relationship is compared to the preferred age difference *D*_*p*_ (which defaults to zero). If the *α*_gap,agescale_ values are set to zero, then *D*_*p*_ will always be the preferred age gap; a positive value between zero and one will cause the preferred age difference to increase as the individual grows older. The *α*_gap,factor_ values at the start of these terms in turn determine the importance of the preferred age difference in the hazard. By setting the *α*_gap,factor_ to a negative value, the hazard decreases as the actual age difference between the candidate partners deviates from the preferred age difference.

The *β* parameter can be used to introduce an effect on the hazard that depends on the time since the relationship became possible. Here, *t*_ref_ refers to the point in time at which the relationship between the two candidate partners became possible. If no relationship existed between them earlier, this is the time at which the youngest person reached the debut age. On the other hand, if they were previously in a relationship with each other already, it is the time at which that relationship was dissolved. For completeness, the factor *F* is a normalization-like factor, to be able to use similar parameters for different population sizes.

A formation event results in the establishment of a sexual relationship, and subsequently, the female partner is at risk of falling pregnant. In that case a *conception event* will be triggered and a while later a *birth event* will take place, introducing a new individual into the population. In case one of the partners in the relationship is HIV-infected, transmission of the virus may occur. If so, a *transmission event* will fire, and the newly infected individual will go through the different infection stages as described earlier. Of course, it is also possible that the relationship will cease to exist, in which case a *dissolution event* will be triggered. Note that in the version at the time of writing, there is no mother-to-child-transmission (MTCT).

Starting ART and dropping out of treatment is managed by another sequence of events. When an individual becomes HIV-infected, either by HIV seeding or by transmission, first a *diagnosis event* is scheduled. Upon diagnosis, an *HIV monitoring event* is scheduled to monitor the progression of the HIV infection. When this event is fired, ART may be initiated, but only if the individual is both eligible (according to a CD4 cell count threshold) and willing to start HIV treatment; if not, a new monitoring event will be scheduled. If ART is initiated, no more monitoring events will be scheduled, but the individual will be at risk of discontinuing his or her HIV treatment, in which case a *dropout event* is triggered. When a person drops out of treatment, a new *diagnosis event* will be scheduled, which should be interpreted as an act of re-engagement in HIV Care^31^.

## Formulating, running and analysing IBMs with SimpactCyan from R or Python

Instructions for installing the core SimpactCyan program and its R interface (the Python interface is automatically installed along with the core program) can be found at http://www.simpact.org/how-to-use-simpact/. To set up a simulation, one needs to prepare a configuration file as a text file with key/value pairs, describing all parameters of the simulation, a snippet of which could look like this:

~~~
…
population.nummen = 200
population.numwomen = 200
population.simtime = 40
…
~~~

Preparing the configuration file manually is time-consuming work however, as *all* event properties needed in a simulation must be set. To make it easier to prepare and run simulations, there is a Python module that can be used to control SimpactCyan from Python, or alternatively an R library that can be installed in R, with a similar interface. It is also possible to use a combined approach: first prepare a configuration file from within R or Python, and subsequently use this configuration to start simulations from the command-line. It can be very helpful when running simulations on a high performance computing cluster for example, where R or Python may not be available.

To use SimpactCyan from within an R session, the `RSimpactCyan` library must first be installed and loaded. This provides a `simpact.run` function that allows a simulation to be configured much more easily than using the configuration file mentioned above: instead of needing to set all parameters of a simulation, only the parameters that are different from the default values need to be specified. The full documentation of all the parameters that can be configured, what they mean and what their default values are, is found at https://simpactcyan.readthedocs.io/en/latest/simpact_simulationdetails.html. If only the key/value pairs in the code snippet above deviate from their default values, the configuration of the simulation would simply become:

~~~
cfg <- list()
cfg[“population.nummen”] <- 200
cfg[“population.numwomen”] <- 200
cfg[“population.simtime”] <- 40
~~~

Similarly, the Python module `pysimpactcyan` defines a `PySimpactCyan` class with a `run` member function that also needs only the settings that differ from the defaults:

~~~
cfg = { }
cfg[“population.nummen”] = 200
cfg[“population.numwomen”] = 200
cfg[“population.simtime”] = 40
~~~

Many of the configuration values will be character strings or numbers, but for some options it is allowed to specify one of the supported one- or two-dimensional probability distributions. For example, the `birth.pregnancyduration.dist.type` is by default set to `fixed` with a value corresponding to 268*/*365 (simulation times are expressed in years), such that every pregnant woman would give birth after precisely 268 days. To allow for some variability (e.g. a standard deviation of 16 days), a log-normal distribution could be used instead:

~~~
mu <- 268/365
var <- (16/365)^2
cfg[“birth.pregnancyduration.dist.type”] <- “lognormal”
cfg[“birth.pregnancyduration.dist.lognormal.zeta”] <- log(mu/sqrt(1+var/mu^2))
cfg[“birth.pregnancyduration.dist.lognormal.sigma”] <- sqrt(log(1+var/mu^2))
~~~

Apart from using a fixed number, supported one-dimensional distributions are the beta, exponential, gamma, log-normal, normal and uniform distributions, as well as user-defined discrete distributions (e.g. based on the frequencies listed in a CSV file). For two-dimensional distributions, one can specify a fixed pair of values, or choose values from binormal or uniform distributions. Here too, user-defined discrete distributions can be specified.

## Runtime analysis

We conducted a small simulation study with SimpactCyan to explore how the runtime varies as a function of population size (at the outset of the simulation), single-core versus multi-core execution of the simulation, fraction of the population of the opposite sex that each individual can potentially form relationships with, and number of relationships that are formed over the course of the simulation. Specifically, we designed scenarios of HIV epidemics in heterosexual populations and ran each scenario over a calendar time period of 40 years. Each scenario was repeated 10 times with different seeds for the random number generators, and the average runtime was calculated per scenario. Scenarios varied by initial population size (4 000, 10 000 and 20 000), the fraction of the population of the opposite sex that each individual could form relationships with (0.2 and 0.4), and the effect of the number of relationships already engaged in by candidate partners on the log-transformed hazard of these two individuals forming a relationship (i.e. the *α*_numrel,man_ and *α*_numrel,woman_ parameters in the hazard function (−0.5 and −5)). These parameters are strongly correlated with the number of relationships that are formed over the course of a simulation. Lastly, we executed simulations with the single-core and multi-core versions of the algorithm. Simulations were run on a MacBook Pro with Intel Core i9 6-core processor, running macOS Mojave (Version 10.14.3).

The results of this simulation study are summarised in Figure 3. Runtimes increase faster than linearly with population size. This is to be expected, since the number of potential relationships that needs to be scheduled in a population of x men and y women is x*y. I.e. this number increases quadratically with population size. Reducing the number of potential partners by setting the `population.eyecap.fraction` parameter to a value smaller than 1, leads to an appreciable reduction in runtime, without affecting the rate at which relationships are formed by much. Running the simulation in parallel (all 6 cores of the machine’s processor are used in the calculations) only leads to a (modest) speedup for scenarios of highly connected networks (many relationships are being formed) in large populations. Typically, simulation studies require several dozens or hundreds of simulation runs, and in that case it is more efficient to distribute single-core simulations in parallel over multiple cores than running multi-core simulations sequentially.

**Figure 3.**
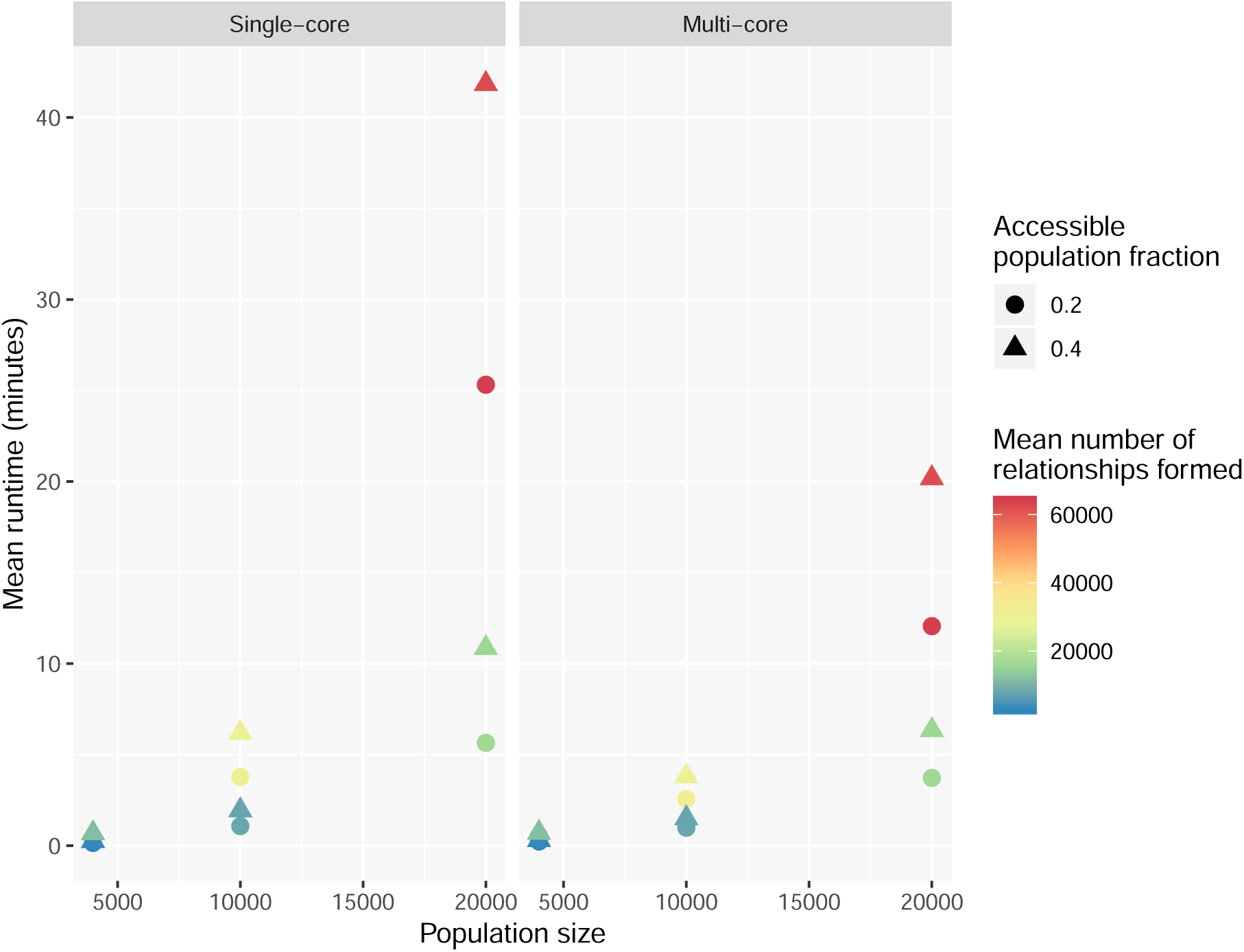
Mean runtime, in minutes, of simulation runs for model scenarios varying in population size, opposite sex population fraction accessible for relationship formation, mean number of relationships formed over the course of the simulation, and computing cores used for each of the runs.

## Model applications

The following section discusses two example simulations that were done using SimpactCyan. The first illustrates how SimpactCyan can be used to assess the impact of progressive changes to the ART eligibility criteria in Eswatini (formerly known as Swaziland). The second illustrates the use of SimpactCyan as a data-generating tool for assessing the performance of other modelling frameworks. All code and data files necessary to reproduce the examples are available at https://github.com/wdelva/SimpactCyanExamples.

### The impact of Early Access to ART for All on HIV incidence

In the MaxART project^32^, SimpactCyan is used to estimate the likely impact of Eswatini’s shift towards “Early Access to ART for All” (EAAA) on the incidence of HIV. HIV incidence is the rate at which HIV-uninfected people acquire the infection. Such infection events are scheduled each time a relationship is formed between an HIV-infected and an HIV-uninfected individual. The hazard for the event is given by

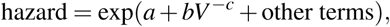

where the other terms are not enabled by default, but allow for a hazard-lowering effect of multiple ongoing relationships (so-called coital dilution^33, 34^), as well as a hazard-increasing effect of adolescent age among women^35^. In this formula, *a, b* and *c* are model parameters; the *V* value represents the current HIV viral load of the person that is already infected.

The viral load model is based upon the notion that an infected person has a specific set-point viral load, *V*_sp_, which corresponds to the viral load in the chronic stage of the infection. The three parameters `person.vsp.toacute.x`, `person.vsp.toaids.x` and `person.vsp.tofinalaids.x` determine the factors by which the HIV transmission hazard should be multiplied during the initial acute stage, as well as the early and late AIDS stages. The *V* value in this hazard expression can therefore be different from the *V*_sp_ value, depending on the time since infection. The non-linear form of this hazard function was inspired by equation (9) published by Hargrove et al.^36^, while the default parameter values are based on a fit to model output generated by Fraser et al.^37^.

At the time of HIV acquisition, time till HIV-related death is determined, based on a paper by Arnaout et al.^38^:

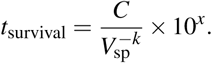

In this formula, *C* and *k* are parameters that can be configured by the user if desired; the *x* parameter (which defaults to zero) is person-specific, and its distribution can be configured to control the amount of variation in post-HIV infection survival times among people with the same set-point viral load.

The set-point viral load value allocated to a newly infected individual is partly determined by that of their infector, i.e. some heritability of set-point viral load is assumed^39^. This is done by using a two-dimensional distribution

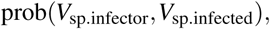

of which the parameters can be chosen using the configuration values. This is subsequently used to obtain the conditional probability when fixing the initial *V*_sp_ value for a person that becomes infected due to the transmission event. By default, a symmetric, truncated bivariate normal distribution with mean 4, minimum 1, maximum 8, standard deviation 1 and correlation coefficient 0.33 is used to sample a log_10_ set-point viral load value for a newly infected person, conditional on the set-point viral load of the infector. To choose the initial set-point viral loads for ‘seed infections’, a marginal probability distribution is used, however.

ART initiation affects both the expected time till HIV-related death and the infectiousness of the person on ART. As soon as ART is started, the log_10_ viral load is assumed to drop by a user-defined fraction, and the updated current viral load is used to re-calculate *t*_survival_. In the EAAA simulation study we assumed that upon ART initiation, the log_10_ viral load drops by 70%, effectively rendering the viral load “undetectable” for most ART clients. Via intervention events, most model parameters can be changed at arbitrary points in time during the simulation. However, person-specific parameter values (e.g. the probability of accepting ART if ART-eligible) and some event-times (e.g. time of non-HIV-related death) are determined at the time the individual is introduced into the population (at the start of the simulation or at birth). Hence, changing related parameters through an intervention event would only affect individuals born into the population after this intervention event, and not the extant population.

In this study, intervention events allowed us to assume that ART was gradually introduced around the year 2000, and that the CD4 cell count threshold for ART eligibility progressively shifted towards ever more inclusive criteria, alongside a decreasing lagtime between HIV infection and HIV diagnosis. These assumptions hold in both the “Status Quo” scenario and the “Early Access to ART for All” (EAAA) scenario. In the EAAA scenario, however, an additional policy change is modelled: a policy of immediate access to ART for all people infected with HIV is adopted from October 2016. In the alternative scenario, the CD4 cell count threshold for ART eligibility stays at 500 cells/microliter from mid 2013 onwards.

The EAAA model was calibrated to demographic, epidemiological and programmatic data (which we refer to as target features) from Eswatini. Specifically, we used the UNAIDS annual national HIV prevalence estimates (1990-2017) and ART coverage estimates (2010-2017)^40^, the estimated average population growth rate over the period 2000 to 2016^41^, the gender- and age group-specific HIV prevalence and incidence estimates from the 2011-2012 SHIMS I study^42^ and the UNAIDS 2017 estimate for the fraction of people aged 15 and above who were virally suppressed (less than 1000 viral copies per mL blood)^40^. Nineteen model parameters were calibrated to these data. Together these parameters determine the sexual behavioural and demographic dynamics of the model population, as well as temporal changes in the rate at which HIV-positive people in the model population were diagnosed with HIV infection, and the extent to which adolescent girls and young women are biologically more susceptible to HIV acquisition than older women and men. In doing so, these 19 parameters drive the model’s features (i.e. summary statistics) that needed to be matched to the target features. Model calibration was achieved by applying the adaptive population Monte Carlo Approximate Bayesian Computation scheme described by Lenormand et al.^43^. Iteratively sampling from the parameter space, starting from the prior distributions (wide-ranged uniform distributions) of the 19 model parameters, the method searches for areas in parameter space that produce model features close to the target features. After thirteen waves of simulations, totalling 29 500 model runs, the convergence criterion was reached and the calibration scheme produced a posterior distribution for the 19 parameters, by way of the 250 best fitting models. Here, “model” means a unique parameter combination producing model features similar to the target features. For each of the 250 models that jointly comprised the parameter posterior distribution, we calculated the root means squared relative error between model features and target features, as a summary measure of goodness-of-fit. The 3 models that fit the data best, which one can think of as the estimated mode of the posterior, were used in the forward projecting step of the analysis. In Supplementary Table S1, we provide the complete list of the 66 summary statistics (target features) that were used to calibrate the model, as well as the corresponding model features, obtained by averaging over the 10 model runs.

In this second part of the analysis, we simulated two scenarios for the expansion of ART in Eswatini, and for each of the two scenarios we ran each of the 3 models 10 times by keeping the model parameter fixed and only changing the seed of the random number generator. In the EAAA scenario, we simply used the 3 best-fitting models and ran them until 2032. In the counterfactual scenario, the same model parameters were used with the exception that the CD4 threshold for ART eligibility remained capped at 500 CD4 cells per microliter from mid 2013 onwards. In all of the subplots of Figure 4, the output from the 3 models is grouped by colour: shades of red for the EAAA scenario and shades of blue for the counterfactual scenario. For each model, the output of the 10 individual model runs is shown in thin dashed lines, and a solid line shows the average model trend. A darker, thick line represents the average across the 30 (3 times 10) runs for each scenario. Grey boxes represent the ranges around the UNAIDS estimates within which the actual numbers lie, based on the best available information^40^. The black dot in Figure 4b indicates the 2017 UNAIDS estimate for HIV incidence among 15-49 year-old adults in Eswatini^44^ (not used for model calibration).

**Figure 4.**
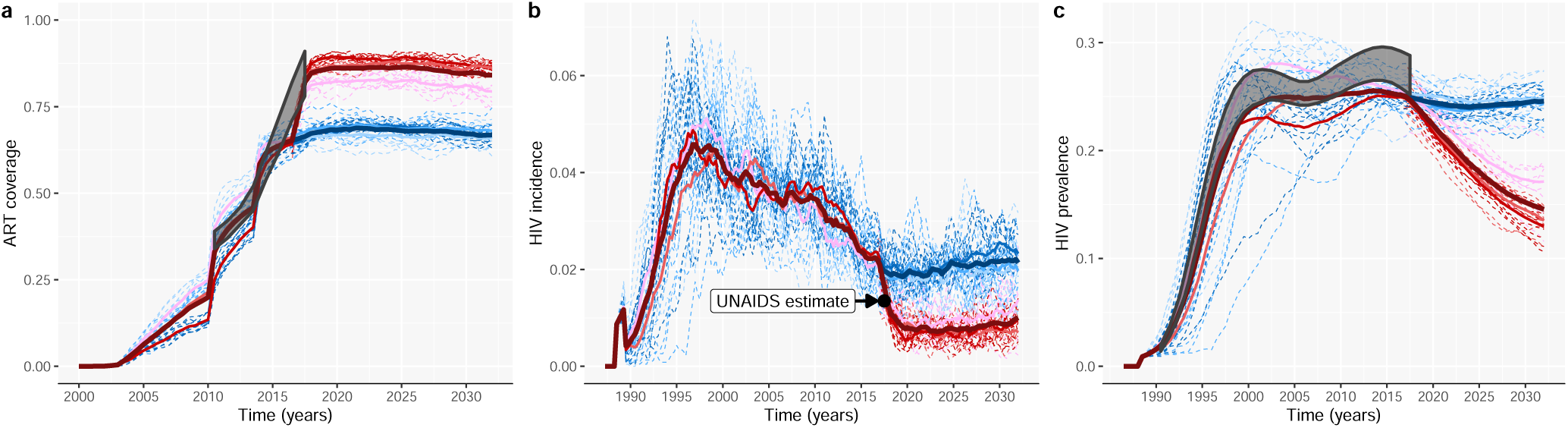
Programmatic and epidemic projections under a “Status Quo” (blue) and “Early Access to ART for All” (red) scenario for the roll-out of a nation-wide ART programme. (**a**) ART coverage: Fraction of the adult HIV-positive population (≥ 15 years old) on ART. (**b**) HIV incidence rate among 15 to 49 year-old people. (**c**) HIV prevalence among 15 to 49 year-old people.

The impact of the policy shift to EAAA was estimated by the relative reduction in HIV incidence (1 minus the ratio of the incidence rates under the two scenarios). Under the counterfactual scenario, HIV incidence was projected to drop by 12% by the end of 2019, from its base level of 2.2 / 100 PY in October 2016. However, under the factual scenario, the models, on average, estimated that the incidence will decrease by 64% over that same period, to 0.8 / 100 PY. The impact of EAAA, as measured by the incidence rate ratio for the 2 scenarios is projected to remain stable over the next decade. The variation between the output of these three models, as shown in Figure 4b, provides a sense of the uncertainty around our best guess of the future impact of EAAA on HIV incidence. It should, however, not be interpreted as an estimate of the credibility or confidence interval, because only the mode of the posterior was used in the impact estimation.

### SimpactCyan as a data-generating benchmarking tool

Phylogenetic trees - branching diagrams depicting the evolutionary relationships among various biological species based on how similar (parts of) their genomes are, have been used as input into so-called phylodynamic models to infer properties of HIV epidemics such as time-trends in HIV incidence rates^45, 46^ or the age-mixing pattern in HIV transmission clusters^47^. Yet, validating these novel modelling frameworks is challenging, given that the true epidemic properties are typically unknown. Likewise, when precise and accurate empirical evidence is lacking, it is difficult to assess to what extent this model-based inference is sensitive to breaches in the model’s assumptions. For instance, the validity of some phylodynamic models hinges on HIV sequence data being available for the majority of HIV-positive people^48^. In most countries in sub-Saharan Africa, however, this is not the case. In Figure 5 we illustrate the use of SimpactCyan to assess the effect of reduced HIV sequence coverage on the correlation between the timing of the internal nodes of the time-resolved phylogenetic tree and the timing of the actual HIV transmission events.

**Figure 5.**
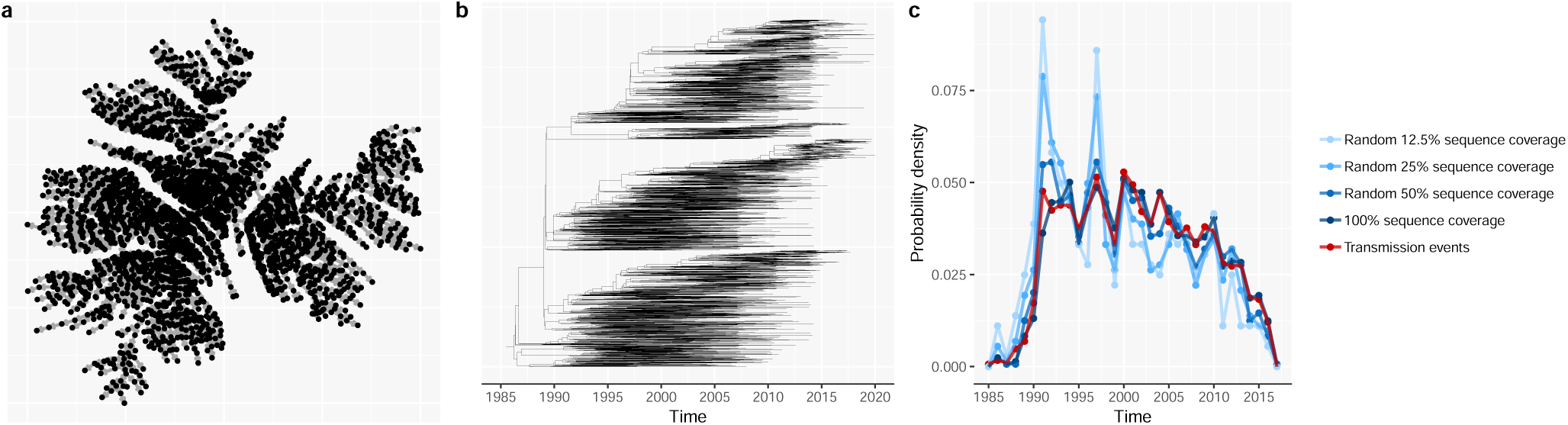
(**a**) The cumulative HIV transmission network, linking all individuals whose HIV infections originated from one seed infector (man 858). (**b**) The time-resolved phylogenetic tree, reconstructed from synthetic HIV sequence data, generated by simulating the molecular evolution of HIV viral strains across the HIV transmission network. (**c**) The probability density of internal nodes in the reconstructed phylogenetic tree(shades of blue) and simulated HIV transmission events (red). The correlation between the timing of the internal nodes and HIV transmission events becomes weaker as the HIV sequence coverage decreases.

First, we simulated an HIV epidemic similar to the epidemics in the EAAA analysis explained in the previous model application. Figure 5a shows the cumulative HIV transmission network that originated from one of the 20 seed infectors (man 858). This network links all individuals who got infected with HIV by man 858 or one of his descendants by the end of the simulation. Next, we converted the transmission network into a phylogenetic tree by assuming that HIV transmission events correspond to branching points in the phylogeny. While this assumption is obviously overly simplistic, it is not an uncommon assumption in phylodynamic modelling studies^49–52^. More importantly, it does not invalidate the didactic purpose of this model application. That purpose is to illustrate SimpactCyan’s potential use in benchmarking the performance of another modelling framework by constructing a range of scenarios in which the extent to which the assumptions made by the other framework are in line with the processes that generated the simulated data. Having converted the transmission network into a phylogenetic tree, we simulated the viral evolution along this phylogeny, using the Seq-Gen program^53^. For this, we assumed a generalised time-reversible substitution model^54^ with a gamma-invariable mixture model (GTR+*γ*+I) for rate heterogeneity among sites. In this way, we generated synthetic HIV sequence data. Specifically, for the root sequence we chose a consensus sequence of the pol region of HIV-1 subtype C, isolated in 1989 from South Africa and retrieved from the HIV Sequence Database of the Los Alamos National Library (LANL) (access number HIV.1.C.ZA.PolCDS1989)^55^. The program jModelTest version 2.1.3^56^ was used for selecting the best fitting evolutionary model to explain the viral diversity in a dataset of HIV-1 subtype C sequences from 386 South African patients, also extracted from the LANL HIV Sequence Database. Using hierarchical likelihood ratio tests, the GTR+*γ*+I model was ranked as the best fitting evolutionary model for these viral sequence data, and the model’s parameter values as estimated by jModelTest were used in the forward viral evolution simulation. Specifically, the relative frequencies of adenine (A), cytosine (C), guanine (G), and thymine (T) were estimated at 0.3906, 0.1752, 0.2201, and 0.2142 respectively. Further, the inferred value for the shape parameter for the *γ* rate heterogeneity was 0.625, and the heterogeneity in transition rate across sites was discretized into 4 discrete *γ* rate categories. The 6 substitution rate parameters of the GTR model were estimated at 1.9803, 9.4404, 0.9423, 0.8770, 11.6367, and 1.0000, with an assumed evolutionary rate of 0.00475 substitutions per site per year (branch length scaling factor)^57, 58^. The fraction of invariant sites was estimated at 0.213. Next, we fed these synthetic sequences into the phangorn^59^ and treedater R packages^60^ to reconstruct the time-resolved phylogenetic tree (Figure 5b), by fitting the GTR+*γ*+I model with a likelihood-based approach and root-to-trip regression. Lastly, we summarised the timing of the internal nodes in the reconstructed time-resolved tree by a vector of the number of internal nodes (i.e. branching points) per calendar year time interval.

In Supplementary Table S2, we show the model parameters of the generalised time-reversible substitution model with a gamma-invariable mixture model (GTR+*γ*+I) for rate heterogeneity among sites, fitted to the empirical sequence data (sample size = 386), and fitted to the simulated sequences (sample size = 2896). To allow for a more direct comparison, we extracted from the complete synthetic sequence dataset a matching sample of simulated sequences with sampling dates as close as possible to those of the empirical sequence data. In Supplementary Figure S1, we also show the empirical phylogenetic tree, the matching sample simulated tree, and a violin plot of the density of patristic distances of the respective trees. Lastly, we report in Supplementary Table S3 nine topological properties of the respective trees. Taken together, this additional information shows strong agreement between empirical and simulated data.

In the ideal case, the sequence dataset includes sequences for all people who were ever infected with HIV, and in addition to that, the molecular evolution model used to reconstruct the phylogenetic tree is exactly the same model that was used to generate the sequence data. Under such ideal circumstances, the timing of the simulated HIV transmission events should correlate strongly with the timing of the internal nodes in the reconstructed tree. Indeed, in a perfect scenario of internal consistency and complete data, the distribution of internal nodes in the reconstructed phylogenetic tree and simulated HIV transmission events matched nearly perfected (red and dark blue lines in Figure 5c). While an exhaustive sensitivity analysis of how phylodynamic inference could be affected by missing data and assumptions that are not consistent with the data-generating processes is beyond the scope of this paper, we simulated three additional scenarios, to illustrate how reduced sequences coverage (50%, 25% and 12.5%) could add noise and bias to the phylodynamic inference. Coverage here is defined as the fraction of people in the cumulative HIV transmission network for whom a consensus sequence is included in the HIV sequence database. In all three of these imperfect scenarios, we still used the appropriate molecular evolution model to reconstruct the phylogenetic tree. As sequence coverage decreases, the timing of internal nodes becomes a less accurate proxy for the timing of transmission events, and hence, a less reliable source for inferring time trends in HIV incidence.

## Future directions

While the present version of SimpactCyan allows users to build models that vary greatly in complexity and application area, there are several ongoing developments that will enable investigations in additional domains. Epidemics of Herpes Simplex Virus 2 (HSV-2) and Hepatitis C Virus (HCV) often emerge as a result of complex interactions with the behavioural and biological factors that drive HIV epidemics. Hence, future versions of SimpactCyan should make joint modelling of these sexually transmitted infections possible. Moreover, studies of HIV transmission, treatment and prevention in injecting drug users (IDU) and children will require additional events for parenteral and mother-to-child transmission of HIV. Lastly, we are pursuing software extensions that will enable investigation of the hypothesis that there are negative correlations between sexual risk behaviours and health seeking behaviours. Indeed, Huerga and colleagues have reported a multivariable analysis in which inconsistent condom use in the preceding year was independently associated with unawareness of being HIV-positive. The same team also reported another multivariable analysis in people aware of their HIV-positive status in which, after adjusting for ARV drug presence in blood, age and sex, individuals with more than one sexual partner had a two times increased risk of being virally unsuppressed^61^.

SimpactCyan was conceived in the spirit of the open science movement^62^ as a flexible open-source, open access tool. We therefore hope that many groups in the field of HIV epidemiology will find it a useful research instrument, and we welcome various applications by others, be it as a data-generating and/or benchmarking tool in methodological research, or for educational purposes. We also believe that the core engine behind SimpactCyan could be used as the starting point for other simulation engines with use cases outside of HIV epidemiology and even outside of epidemiology altogether.

## Acknowledgements

The authors thank Jonathan Dushoff and Roxanne Beauclair for valuable comments on earlier drafts of this manuscript. The computational resources and services used for the calibration of the EAAA model were provided by the VSC (Flemish Supercomputer Center).

## Funding sources

This work was supported by grants G091210N, G0B4314N and W002514N from the Research Foundation – Flanders (FWO), grant ZEIN2010PR375 from the Flemish Interuniversity Council (VLIR), grant 100014 from NRF-TWAS, and a grant from the Dutch Postcode Lottery.

## Author contributions

JL wrote all the source code of the core engine and the R and Python interfaces presented in this paper, and wrote the first draft of the manuscript. DMH drafted the introduction section and Figure 2. EK contributed to Figure 2 and provided editorial assistance. DN wrote R code for the second example application. NH contributed to the software design and provided editorial assistance. WD contributed to the software design, performed the runtime analysis, wrote R code for the example applications, and wrote the “Model applications” and “Future directions” sections of the manuscript. All authors reviewed the manuscript.

## Competing interests

The authors declare no competing interests.

## Notes

#### Summary of Updates

Minor textual edits in introduction and discussion section.

https://github.com/wdelva/SimpactCyanExamples

