## Supplementary Tables S1, S2, S3 and Figure S1 for "SimpactCyan 1.0: An Open-source Simulator for Individual-Based Models in HIV Epidemiology with R and Python Interfaces"

**Table S1. Model calibration results: model and target features.**

| | Model 1 | Model 2 | Model 3 | Target feature | Source |
|---|---|---|---|---|---|
| ***Demography*** | | | | | World Bank, 2018 |
| Average annual population growth rate in 2000 - 2016 | -0.20% | 1.10% | 2.20% | 1.50% | |
| ***HIV prevalence*** | | | | | Bicego et al. 2013 |
| 18-19 year-old women | 18.30% | 17.00% | 23.10% | 14.30% | |
| 18-19 year-old men | 1.40% | 1.20% | 3.40% | 0.80% | |
| 20-24 year-old women | 29.20% | 30.10% | 32.80% | 31.50% | |
| 20-24 year-old men | 7.50% | 7.90% | 10.30% | 6.60% | |
| 25-29 year-old women | 47.00% | 44.80% | 41.90% | 46.70% | |
| 25-29 year-old men | 21.40% | 21.70% | 23.80% | 21.30% | |
| 30-34 year-old women | 52.70% | 55.50% | 44.60% | 53.80% | |
| 30-34 year-old men | 35.50% | 36.90% | 38.30% | 36.60% | |
| 35-39 year-old women | 50.80% | 50.80% | 41.80% | 49.10% | |
| 35-39 year-old men | 38.50% | 42.40% | 42.50% | 47.00% | |
| 40-44 year-old women | 45.00% | 46.70% | 42.40% | 39.70% | |
| 40-44 year-old men | 37.90% | 41.80% | 41.70% | 45.50% | |
| 45-49 year-old women | 39.70% | 43.00% | 33.40% | 31.60% | |
| 45-49 year-old men | 33.60% | 36.40% | 32.40% | 42.50% | |
| ***HIV incidence (per 100 person-years)*** | | | | | Justman et al. 2017 |
| 18-19 year-old women | 6.3 | 5.2 | 6 | 3.8 | |
| 18-19 year-old men | 0.8 | 1 | 1.9 | 0.8 | |
| 20-24 year-old women | 5.2 | 5 | 4.7 | 4.3 | |
| 20-24 year-old men | 2.6 | 2.8 | 2.9 | 1.6 | |
| 25-29 year-old women | 2.4 | 3.2 | 2.8 | 2 | |
| 25-29 year-old men | 4.7 | 4.3 | 4.8 | 2.6 | |
| 30-34 year-old women | 1.5 | 5.6 | 2.2 | 2.7 | |
| 30-34 year-old men | 2.9 | 4.7 | 5.4 | 3.1 | |
| 35-39 year-old women | 1.3 | 0.7 | 0.9 | 4 | |
| 35-39 year-old men | 2.5 | 3.4 | 2.5 | 0.4 | |
| 40-44 year-old women | 0.8 | 0.2 | 0.6 | 2.1 | |
| 40-44 year-old men | 1.4 | 1.7 | 2.8 | 1.2 | |
| 45-49 year-old women | 0 | 0.7 | 0.3 | 1.2 | |
| 45-49 year-old men | 0 | 1.4 | 1.3 | 0 | |
| ***ART coverage in >=15 year-old adults*** | | | | | UNAIDS, 2019 |
| 2010 | 39.90% | 34.20% | 25.00% | 37% | |

| | | | | | |
|---|---|---|---|---|---|
| 2011 | 46.10% | 40.30% | 32.00% | 40% | |
| 2012 | 49.40% | 43.60% | 36.60% | 44% | |
| 2013 | 51.30% | 46.00% | 40.20% | 49% | |
| 2014 | 64.30% | 62.20% | 58.50% | 58% | |
| 2015 | 64.80% | 64.30% | 62.10% | 67% | |
| 2016 | 65.10% | 66.20% | 64.20% | 76% | |
| 2017 | 78.90% | 83.10% | 84.20% | 85% | |
| ***Virology*** | | | | | UNAIDS, 2019 |
| fraction of >=15 year-old adults who were virally suppressed (less than 1000 viral copies per mL blood) | 82.50% | 85.80% | 86.80% | 74% | |
| ***HIV prevalence in 15-49 year-old adults*** | | | | | UNAIDS, 2019 |
| 1990 | 2.20% | 1.70% | 1.90% | 1.70% | |
| 1991 | 3.90% | 2.40% | 3.10% | 3.30% | |
| 1992 | 6.10% | 3.80% | 5.00% | 5.70% | |
| 1993 | 8.60% | 5.60% | 7.90% | 8.80% | |
| 1994 | 11.80% | 7.90% | 11.50% | 12.40% | |
| 1995 | 15.40% | 11.10% | 15.40% | 16.10% | |
| 1996 | 18.90% | 13.90% | 18.60% | 19.40% | |
| 1997 | 21.90% | 16.40% | 20.60% | 22.00% | |
| 1998 | 24.50% | 18.90% | 22.00% | 23.90% | |
| 1999 | 26.20% | 20.80% | 22.60% | 25.10% | |
| 2000 | 27.10% | 22.00% | 22.90% | 25.80% | |
| 2001 | 27.60% | 22.80% | 23.00% | 26.10% | |
| 2002 | 28.10% | 23.50% | 22.90% | 26.10% | |
| 2003 | 28.10% | 24.20% | 22.50% | 25.90% | |
| 2004 | 27.90% | 24.50% | 22.40% | 25.70% | |
| 2005 | 27.70% | 24.80% | 22.30% | 25.50% | |
| 2006 | 27.40% | 24.80% | 22.20% | 25.60% | |
| 2007 | 27.30% | 24.80% | 22.60% | 25.90% | |
| 2008 | 26.80% | 25.00% | 22.80% | 26.30% | |
| 2009 | 26.90% | 25.30% | 23.20% | 26.80% | |
| 2010 | 26.70% | 25.40% | 23.70% | 27.40% | |
| 2011 | 26.30% | 25.40% | 24.20% | 27.80% | |
| 2012 | 26.10% | 25.40% | 24.60% | 28.20% | |
| 2013 | 26.20% | 25.50% | 25.00% | 28.40% | |
| 2014 | 25.90% | 25.50% | 25.00% | 28.40% | |
| 2015 | 25.70% | 25.20% | 25.00% | 28.30% | |
| 2016 | 25.50% | 25.00% | 24.90% | 27.90% | |
| 2017 | 25.10% | 24.40% | 24.50% | 27.40% | |

**Table S2. Evolutionary models fitted to empirical and synthetic data.**

| | Empirical data | Synthetic data |
|---|---|---|
| ***Relative Frequencies*** | | |
| adenine (A) | 0.3906 | 0.3929 |
| cytosine (C) | 0.1752 | 0.1726 |
| guanine (G) | 0.2201 | 0.2234 |
| thymine (T) | 0.2142 | 0.2111 |
| ***Rate heterogeneity*** | | |
| shape parameter | 0.6250 | 0.6244 |
| ***Relative substitution rates*** | | |
| r(A → G) = r(G → A) | 1.9803 | 2.0421 |
| r(A → C) = r(C → A) | 9.4404 | 9.5318 |
| r(A → T) = r(T → A) | 0.9423 | 0.9674 |
| r(G → C) = r(C → G) | 0.8770 | 0.8840 |
| r(G → T) = r(T → G) | 11.6367 | 12.0604 |
| r(C → T) = r(T → C) | 1.0000 | 1.0000 |
| ***Fraction of invariant sites*** | | |
| I parameter | 0.2130 | 0.3091 |

**Table S3. Normalised topological properties of the phylogenetic trees reconstructed from empirical and synthetic data with matching sampling dates (See also Figure S1).**

| | Empirical data | Synthetic data |
|---|---|---|
| Sackin index of tree imbalance | 0.1020 | 0.0880 |
| Colless index of tree imbalance | 0.0760 | 0.0590 |
| Average size of ladders[1] | 0.0060 | 0.0070 |
| Cherries[2] | 0.6630 | 0.6350 |
| IL number[3] | 0.3390 | 0.3660 |
| Maximum height of the tips | 0.0830 | 0.0730 |
| Pitchforks[4] | 0.4510 | 0.4770 |
| First staircase-ness measure[5] | 0.6000 | 0.6320 |
| Second staircase-ness measure[6] | 0.6030 | 0.5980 |

1 A ladder is defined as a series of consecutive nodes in the tree, each of which has exactly one tip child. The size of the ladder is given by the number of nodes in the chain.

2 A cherry is a pair of sister tips.

3 The IL number is defined as the number of internal nodes with a single tip child.

4 Pitchforks are clades with three tips.

5 The proportion of subtrees that are imbalanced (i.e. subtrees where the left child has more tip descendants than the right child, or vice versa).

6 The average of all the min(l,r)/max(l,r) values of each subtree, where l and r are the number of tips in the left and right children of a subtree.

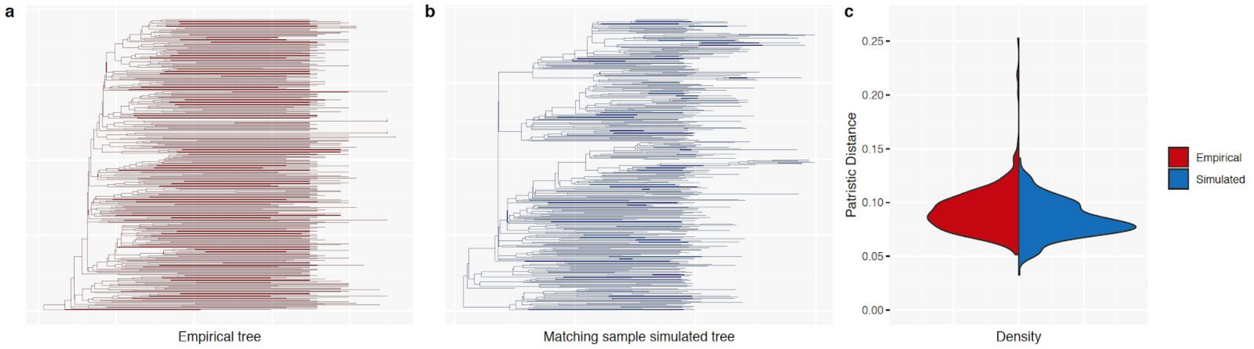


**Figure S1**. (**a**) Time-resolved phylogenetic tree, reconstructed from the empirical HIV sequence data. (**b**) Time-resolved phylogenetic tree, reconstructed from a subset of the synthetic HIV sequence data, with sampling dates that match those of the empirical dataset. (**c**) The density distribution of patristic distances of the respective phylogenetic trees.
